# The DREAM implant: A Lightweight, Modular and Cost-Effective Implant System for Chronic Electrophysiology in Head-fixed and Freely Behaving Mice

**DOI:** 10.1101/2024.02.26.582029

**Authors:** Tim Schröder, Robert Taylor, Muad Abd El Hay, Abdellatif Nemri, Arthur França, Francesco Battaglia, Paul Tiesinga, Marieke L. Schölvinck, Martha N. Havenith

## Abstract

Chronic electrophysiological recordings in rodents have significantly improved our understanding of neuronal dynamics and their behavioral relevance. However, current methods for chronically implanting probes present steep trade-offs between cost, ease of use, size, adaptability and long-term stability.

**SUMMARY:** Introducing a lightweight, cost-effective probe implant system for chronic electrophysiology in rodents, optimized for ease of use, probe recovery, experimental versatility and compatibility with behavior.

This protocol introduces a novel chronic probe implant system for mice called the DREAM (Dynamic, Recoverable, Economical, Adaptable and Modular), designed to overcome the trade-offs associated with currently available options. The system provides a lightweight, modular and cost-effective solution with standardized hardware elements that can be combined and implanted in straightforward steps and explanted safely for recovery and multiple re-use of probes, significantly reducing experimental costs.

The DREAM implant system integrates three hardware modules: (1) a microdrive that can carry all standard silicon probes, allowing experimenters to adjust recording depth across a travel distance of up to 7mm; (2) a 3D-printable, open-source design for a wearable Faraday cage covered in copper mesh for electrical shielding, impact protection and connector placement, and (3) a miniaturized head-fixation system for improved animal welfare and ease of use. The corresponding surgery protocol was optimized for speed (total duration: 2 hours), probe safety and animal welfare.

The resulting implants had minimal impact on animals’ behavioral repertoire, were easily applicable in freely moving and head-fixed contexts and delivered clearly identifiable spike waveforms and healthy neuronal responses for weeks of data collection post-implant. Infections and other surgery complications were extremely rare.

As such, the DREAM implant system is a versatile, cost-effective solution for chronic electrophysiology in mice, enhancing animal well-being, and enabling more ethologically sound experiments. Its design simplifies experimental procedures across various research needs, increasing accessibility of chronic electrophysiology in rodents to a wide range of research labs.

## INTRODUCTION

Electrophysiology with chronically implanted silicon probes has emerged as a powerful technique for investigating neural activity and connectivity in behaving animals, particularly in mice, due to their genetic and experimental tractability^1^. Especially laminar silicon probes have proven to be an invaluable tool to identify functional relationships within cortical columns^2^ and for relating the dynamics of large neuronal populations to behavior in a way that was impossible previously^3^.

Two complementary approaches are the current gold standards for recording neural activity in vivo: two-photon microscopy^4, 5^ and extracellular electrophysiology^6^. The choice of recording methodology constrains the nature of the readouts that can be obtained: two-photon microscopy is particularly well-suited to longitudinal studies of individually identifiable neurons in large populations across time, but suffers from high equipment costs, and is limited to superficial layers of the cortex in intact brains. In addition, the typical temporal resolution of ∼30Hz limits its ability to capture ongoing neuronal dynamics^7, 8^.

In contrast, electrophysiological recordings offer high temporal resolution (up to 40 kHz) to track neuronal activity moment by moment, can be applied widely across species as well as across cortical depths, and setups are relatively low-cost compared to a two-photon microscope. However, identification of individual neurons, as well as longitudinal tracking of neuronal populations are difficult to achieve. This especially applies to wire electrodes, e.g. tetrodes, and to acute electrode insertions. Besides lacking the ability to track neurons across recording sessions^9^, repeated acute insertions cause local trauma^10^ that mounts an immune response^11^, increasing the chance of infection and gliosis. This ultimately reduces the stability of recorded neuronal activity and life expectancy of experimental animals, limiting the scope of longitudinal studies featuring acute electrophysiological recordings to just a few days^12^.

Chronic high-density silicon probe recordings aim to combine some of the best attributes of acute electrophysiology and two-photon imaging, being able to track neural population dynamics across sessions with only somewhat lowered ability to identify individual neurons compared to two-photon imaging^13^, while providing high flexibility in the spatial placement and precise temporal resolution of the recorded signals, as well as improved longevity and wellbeing of experimental animals compared to acute recordings^14^. Furthermore, in contrast to acute recordings, chronic electrophysiology necessitates only a single implantation event, effectively reducing the risk of infection and tissue damage and minimizing stress on the animals^15^. Collectively, these advantages make chronic electrophysiology a powerful tool for investigating the organization and function of the nervous system.

However, commonly used chronic implantation techniques for mice constrain researchers to make significant trade-offs between compatibility with behavioral recordings, implant weight, replicability of implants, financial costs, and overall ease of use. Many implant protocols are not designed to facilitate re-use of probes^16^, steeply raising the effective cost of individual experiments, and thus making it financially difficult for some labs to use chronic electrophysiology. They also often require extensive in-house prototyping and design work, for which the expertise and resources may not be present.

On the other hand, integrated implant systems^17^ offer a more widely accessible solution for chronic electrophysiology in rodents. These systems are designed to integrate a microdrive holding the probe with the remainder of the implant to simplify implant handling and surgical procedures. However, once implanted, such systems can be top-heavy, and limit the experimenter’s ability to flexibly adapt an experiment to different target coordinates. Often, their weight precludes implants in smaller animals, and potentially impairs animal movement and induces stress^18^. This can disproportionately affect research on juvenile and female cohorts, as weight limitations are more likely to affect these groups.

Additionally, not all integrated systems allow for adjustment of electrode positions post implantation. This is relevant, as gliosis or scarring due to probe insertion^19^, especially in the initial 48 hours after implantation^20^, can reduce the quality of the recorded neuronal activity. Micro-adjustments to the probe insertion depth can limit these negative effects on signal integrity. Therefore, micropositioning mechanisms, commonly called microdrives, can be beneficial even in probes with a large number of electrodes distributed across their length.

To overcome such trade-offs, we introduce a novel chronic electrophysiology implant system for mice that addresses the limitations of previous designs by offering a lightweight, cost-effective, and modular solution. The DREAM implant system is designed to weigh less than 10% (approx. 2.1g) of a mouse’s typical body weight, ensuring animal welfare and minimal impact on behavior. Validation of the DREAM implant design shows minimal impact on behavioral key metrics such as locomotion – which can be significantly impacted in rodents when loads are placed on the cranium. This can benefit experimental paradigms that utilize freely moving as well as head-fixed animals, by boosting animal well-being and allowing more ethologically sound experiments.

The system includes a microdrive for flexible adjustment of recording depth up to 7mm and can be adapted to different types of probes and recording devices, providing researchers with a cost effective and versatile tool for various experimental applications. The system is routinely combined with a metal microdrive^21^, which offers consistent probe-recovery compared to other systems (expected average recovery rate: approx. three reliable reuses per probe) and drastically reduces the cost of individual experiments.

The design features a 3D-printed protective Faraday cage, allowing for cheap yet robust protection from electrophysiological noise, mechanical impacts, and infectious materials, enabling stable and noise-free recordings that suffer from minimal infection rates. This implantable cage consists of the so-called ‘crown’, designed for impact protection and to provide structure for the conductive metal mesh coating of the Faraday cage; and the crown ring, which serves as a mount for an implantable amplifier and/or probe connector (see Figure 1).

**Figure 1.**
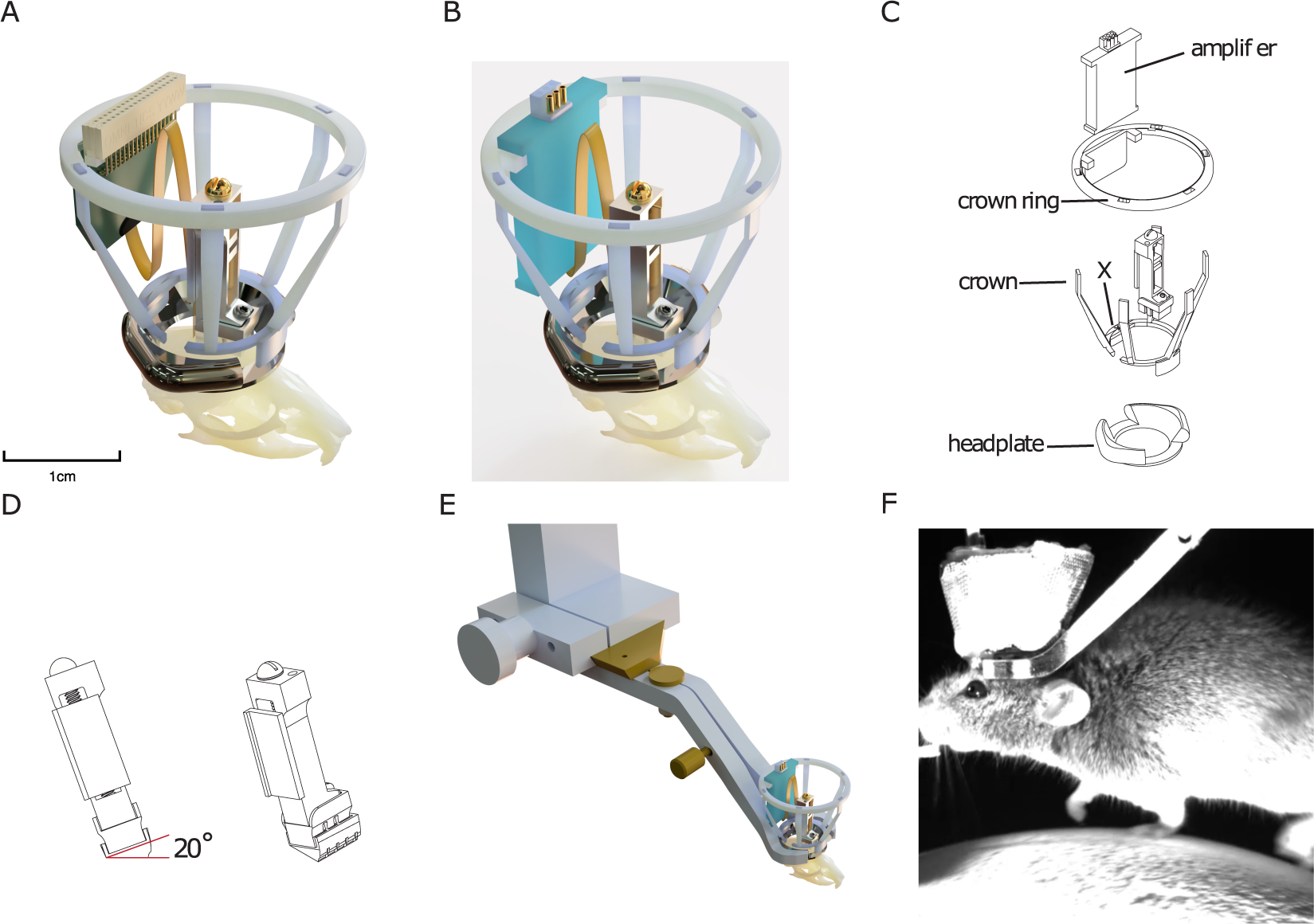
Implant design. (A) 3D rendering of the implant superimposed onto a mouse skull with a silicon probe connected to a probe connector. The central aperture of the headplate is approximately 10mm for scale. The height of the drive is approximately 17mm. The copper mesh that forms the outside of the Faraday crown, as well as ground/ref wires, are not shown. (B) Same as (A) with connection to an amplifier board instead of a probe connector. (C) Exploded technical drawing of the implant, showing its components. (D) Rendering of an angled spacer that can be implanted underneath a microdrive, allowing to consistently implant the microdrive at a predefined angle (here: 20°). (E) Rendering of integrated head-fixation mechanism, showing implanted headplate with Faraday crown with the surrounding head-fixation clamp and the dove-tail connection to setup. (F) Image of mouse head-fixed on a treadmill using the implant’s integrated head fixation mechanism.

Finally, the headplates included in the modular implant system are designed to be compatible with a novel, efficient head-fixation system without adding extra bulk to the implant. In contrast to other existing systems, it does not require tightening small screws close to the implant, speeding up fixation of mice in the experimental setup, improving the experimenter-animal relationship, as well as behavioral adherence. At the same time, the headplate is used as a base on which to build the other modules of the DREAM chronic electrophysiology system.

Design files for the DREAM implant are published as open-source hardware at https://github.com/zero-noise-lab/dream-implant/. In the following sections, the design and fabrication of the DREAM implant system will be described, its successful implementation in a mouse model will be demonstrated, and its potential applications and advantages compared to existing systems will be discussed.

## PROTOCOL

All experimental procedures were conducted according to the institutional guidelines of the Max Planck Society and approved by the local government’s ethical committee (Beratende Ethikkommission nach §15 Tierschutzgesetz, Regierungspräsidium Hessen, Project approval code: F149-2000).

## 1 Pre-surgical preparations

### 1. Preparation of the silicon probe

1.1. In case of probe re-use, clean the silicon probe according to the recommendations of the probe supplier. It is recommended to soak the probe in enzymatic cleaner (see Table of Materials) for 5–10 minutes, then rinsing it in demineralized water. This should be done as quickly as possible after explantation. A day before (re-)implantation, soak the probe in 70% ethanol for at least 30 minutes for disinfection.
1.2. Measure channel impedances to make sure they are within specifications for the recorded signal. Follow the protocol for testing noise levels from the Neuropixels user manual^22^, measure impedance via the desired recording software (e. g. https://open-ephys.github.io/gui-docs/User-Manual/Plugins/Acquisition-Board.html#impedance-testing) and follow the target channel impedances from the silicon probe manufacturer or datasheet. If impedances are too high, consider recoating of the electrode sites^23^.
1.3. Solder a .05” solder tail socket (see Table of Materials) to the ground (GND) wire of the probe. The socket will be connected to the GND pin (next step) during surgery. In this protocol, a separate reference (REF) pin is not used, as GND and REF are shorted on the headstage used. Therefore, only the GND pin will be mentioned in the remainder of the protocol. If a separate REF is used, repeat the following step for the REF pin.
1.4. To prepare the GND pin, repeatedly insert the pin side of a .05” solder tail socket (see Table of Materials) into the GND .05” solder tail socket until insertion is largely effortless. Using gold-plated pins can reduce the need for this smoothing step. This makes sure that the GND pin and socket can easily be connected during surgery without the need to apply excessive pressure, reducing the risk of injuries to the animal and probe damage.
1.5. If an implantable pre-amplifier for the silicon probe is used, prepare them for chronic implantation following the supplier’s procedures. This might include coating them in silicon or epoxy to avoid moisture damaging the electronics, as well as repeatedly mating the amplifier connector to reduce mating force when connecting the amplifier to the recording system during recordings. This is especially useful for Omnetics users. Then attach the amplifier/connector to the ring of the Faraday cage by using silicone plaster to glue it to the area of the Faraday ring designed to hold the amplifier (see Figure 1).

### 2. Preparation of the microdrive and headgear

2.1. Turn the screw on the microdrive body so that the microdrive shuttle is almost entirely retracted upwards.
2.2. Optionally, attach an angled spacer (see Figure 1D) to the bottom of the microdrive with cyanoacrylate glue or dental cement which can be used to allow for a specific degree of tilt to be used, for example when recording through cortical layers in a region within the central sulcus, or within deep structures that may require a non-perpendicular approach (for angled spacer, see Table of Materials).
2.3. Lay the microdrive horizontally onto the microdrive holder. (Supplementary Figure S1)
2.4. Place a small piece of adhesive putty (see Table of Materials) on the microdrive holder, at a distance above the microdrive at which the head-stage connector will be placed. This distance depends on the length of the flex cable that connects the silicon probe to the headstage connector.
2.5. Place a tiny drop of silicone plaster (see Table of Materials) onto the shuttle.
2.6. Take the silicon probe out of its packaging with the help of a blunt soft tipped forceps. These can be made by coating standard needle-nose forceps with 3mm diameter heat-shrink tubing (see Table of Materials). Place the probe with the flex-cable first onto the shuttle of the microdrive, so that the bottom edge of the flex cable hangs slightly over the bottom edge of the microdrive shuttle.
2.7. Gently pull the flex cable towards the top of the microdrive until the bottom edge of the flex cable meets the bottom edge of the microdrive shuttle. Make sure to push the flex cable against the left edge of the microdrive shuttle during this step, so that it is placed exactly vertically on the microdrive in the end. At this point, the shanks of the silicone probe should typically not (or only minimally) protrude past the lower edge of the microdrive (depending on the exact length of the probe shanks and the depth of the targeted brain area).
2.8. Place the head-stage connector of the probe onto the adhesive putty at the top of the holder to protect the probe from falling off.
2.9. Use a 27G syringe needle to apply a small drop of cyanoacrylate glue (see Table of Materials) between the flex-cable and shuttle to secure the probe in place. Very important: Make sure the glue does not run onto the microdrive or along the flex cable beyond the shuttle!
2.10. Once the flex cable is glued in position, attach amplifier to crown ring (see Table of Materials) using silicone plaster, attach the flex-cable to the amplifier and cover the connection and cable in a thin layer of silicone plaster.
2.11. Once 5 minutes has elapsed, and the plaster has set, store the microdrive and probe safely until further use.
2.12. Cut pieces of copper mesh (see Table of Materials) into an open donut shape (see cutting pattern in Supplementary Figure S2) to cover the Faraday cage.
2.13. Fasten the copper mesh cut-out onto the Faraday cage with small drops of epoxy resin (see Table of Materials). For this step one can also replace epoxy with dental cement. The Faraday cage contains a space to house a probe connector or amplifier. This space is marked by an X in the design file, and it contains a supporting base for the amplifier/connector, as well as a larger distance between the two adjacent spokes of the cage. To create sufficient space around the amplifier/connector, fix a small amount of extra mesh between the two adjacent spokes, creating a protrusion. This ensures that the amplifier/connector can later be positioned in this ‘pocket’ without touching the Faraday cage. Note: To ensure secure adhesion with minimal warping, use the crown ring placed directly on the crown to maintain shape and to support the thin spokes of the crown. Furthermore, use soldering helping hands to secure the crown and mesh during drying. If one struggles maintaining the shape of the crown when undergoing the procedure, attempt to only epoxy two of the crown arms at a time to prevent warping.
2.14. If separate grounding of the Faraday cage is desired, solder a small header pin onto a 30mm grounding wire (see Table of Materials), then use conductive epoxy to adhere the wire to the copper mesh cut-out. Note: Our lab does not adhere to this step.

At this point, prepared parts can be safely stored, and surgery can be performed at a later stage.

Implantation of the microdrive and headgear

### 3. Surgery: Preparation of probe and workspace

3.1. Sterilize and place surgical instruments in the surgical workspace following an approved procedure. This can include using a bead sterilizer, autoclaving instruments or rinsing with 30% peroxide or 90% ethanol, depending on the approved experimental protocol.
3.2. Place the ceramic dish used to prepare the dental cement in an ice box, fridge, or freezer following the instructions in the dental cement kit (see Table of Materials). The cooled ceramic dish should be used during cement mixing to increase the time during which the cement is malleable. Use a cooled dish whenever longer cementing steps are required.
3.3. If histological verification of probe placement at the end of the experiment is desired, extend the silicon probe right before the surgery by turning the screw on the microdrive counterclockwise and apply a lipophilic dye (see Table of Materials) to the probe by dipping it in a small drop of the dye. The lipophilic dye can be prepared from a commercially bought DMSO or EtOH diluted stock solution (see Table of Materials) by diluting it in a suitable buffer such as PBS at a 1 to 5 μM concentration.

### 4. Surgery: Preparation of the animal

4.1. Follow an approved anesthesia protocol for a 2-4 hour rodent surgery under aseptic conditions. This can include general and local anesthesia, analgesia, application of eye ointments, and injections of saline. Our lab uses injectable anesthesia (ketamine 100 [mg/kg]/medetomidine 0.5 [mg/kg]) together with local analgesia cream and eye ointment (see Table of Materials), and the animal is placed on a heating pad to regulate body temperature.
4.2. When the animal is fully anaesthetized, move it to a separate non-sterile shaving area. Ensure that the animal is warmed sufficiently, for example by placing it on a heating pad. Remove hair on the top of the skull. This can be done with an electric shaver or depilation cream (see Table of Materials), or by repeatedly shaving the top of the head with a scalpel covered in 70% ethanol. Carefully remove loose hairs to make sure they do not get in contact with exposed tissue later. To remove hairs, use e.g. tissues wetted with 70% ethanol and/or a squeeze ball pump. If using depilation cream, this also needs to be thoroughly removed using cotton swabs and saline.
4.3. Disinfect the shaved area multiple times with an iodine-based disinfectant (see Table of Materials) and alcohol using cotton swabs, moving from the center of the head to the sides to brush any remaining lose hairs away from the incision site.
4.4. Disinfect the fur on and around the head using betadine. This ensures a sterile working area and protects surgical instruments and materials from coming into contact with unsterile fur.
4.5. Place animal in a stereotactic frame using ear bars and nose holder (see Table of Materials).
4.6. Using small surgical scissors (see Table of Materials), cut an almond-shaped opening in the skin on top of the skull, reaching from just posterior of the lambda suture to between the eyes.
4.7. Remove subcutaneous membrane and periosteum, by cutting away while still wet, then scratching the skull with a scalpel blade to remove soft membrane tissue on the surface of the skull that may impede adhesion of dental cement.
4.8. Optional: Once the skull has been cleared of membrane tissue, briefly apply a thin layer of 0.5% peroxide and wash it off with water-based iodine disinfectant (e.g., Betadine) before roughening the surface of the skull to improve adhesion of the primer to the skull.
4.9. Carefully roughen the surface of the skull by scratching a crisscross pattern with the tip of the scalpel turned upside down. This helps dental cement to adhere to the skull later. Note: Do not scratch too vigorously on top of sutures since this can cause the sutures to rupture and leak intracranial fluid, which impairs adhesion of the dental cement.
4.10. Alternate between scalpel blade and sterile cotton buds to gently scratch/push away neck muscles attached to the sides of the lambda suture, until the muscles have been pushed back to the ‘edge’ of the skull on top of the cerebellum. This helps to minimize muscle noise in neuronal recordings.
4.11. Fill an 1ml syringe with a 27G needle (see Table of Materials) with small amounts of surgical cyanoacrylate glue (see Table of Materials). Then glue the skin to the skull edges by using the syringe to smear tiny drops of superglue across it. Glue tissue as flat as possible to the skull to leave space for implants. This procedure ensures that skin and muscles do not come in direct contact with parts of the implant, which avoids muscle noise in recordings, and improves the adhesion of the dental cement.
4.12. Apply dental cement primer across the skull for extra adhesion and harden with UV light (see Table of Materials). This improves dental cement adhesion and prevents cranial sutures from leaking and weakening the cranial-cement bond over time.
4.13. Find the target location for the probe implantation relative to bregma or lambda and draw the outline of the craniotomy around it with a surgical marker. Place the headplate on the skull so that the craniotomy lies within it, with space for the microdrive at one side of the craniotomy, as well as for 1-2 grounding pins.
4.14. Implant the headplate using dental cement. Mix dental cement in the designated cooled ceramic dish (see Step 3.2). Make sure that the headplate adheres to the skull on all sides, forming a watertight ‘well’.
4.15. With a dental drill (size US ½ HP), drill a small burr hole the width of the header pins prepared in Step 1.4 over the brain area(s) to be used as GND/REF. If grounding the Faraday cage is desired, drill another small craniotomy close to the edge of the Faraday cage for the Faraday-GND header pin. Note: For the GND/REF header pin(s), place the craniotomy at a sufficient distance from the edge of the cage that the header pin itself can be placed within it later without touching the Faraday cage.
4.16. Clean the craniotomy by gently dripping sterile saline onto it with a syringe, and removing the saline with non-shedding wipes (see Table of Materials). Repeat until all blood and loose tissue is removed.
4.17. Prepare a 0.7% agar (see Table of Materials) solution in saline, cool slightly and introduce into the craniotomy using a 27G needle on a 1ml syringe.
4.18. Gently insert a GND pin (see step 1.3) into each craniotomy drilled in the previous step. The pin(s) will be surrounded by agar on all sides (see step 4.17). Apply cement around the header pins to secure them and provide electrical isolation.
4.19. Clean the ceramic dish and place it back in the fridge/freezer.
4.20. With a dental drill, drill the outline of a larger craniotomy (circular or square) by moving around the edge in steady movements. The craniotomy should have a size of 1×1mm – 2×2mm to allow for small adjustments to the placement of the probe in order to e.g. avoid blood vessels without exposing too much of the cortex. If possible, avoid placing craniotomies over sutures. Drill in rounds of 20–30 seconds, and cool down the skull with saline between drilling rounds. Note: When beginning drilling, it is useful to mark out the leading edge of the microdrive with a marker, hence ensuring that when drilling, a straight edge can be formed in parallel to the microdrive leading edge. This improves the chances of avoiding cement in the craniotomy when fixing the microdrive in place, as well as improving adhesion, preventing microdrive overhang over the craniotomy and allowing for greater lateral maneuverability when placing the microdrive in relation to final recording site position.
4.21. After a few initial rounds of drilling, test the resistance of the drilled-out portion of bone by gently pushing on it with fine forceps (Size 5 or finer, see Table of Materials). Keep testing in between drilling rounds until the bone begins to ‘bounce’ underneath the forceps when pushed. When this is the case, add a drop of saline on top of the craniotomy to soften up the bone, then use the forceps to gently remove the drilled-out piece of bone. If the bone cannot yet be removed gently, do another round of drilling, focusing on the points where the bone is still attached more strongly. In general, aim to remove the skull with gentle pressure from the forceps before it has been entirely drilled through, since this typically minimizes tissue damage. Note: Ensure the surface of the dura is moistened regularly, both during drilling to reduce temperatures, but also following bone flap removal. This improves the chances of easy probe insertion by preventing the dura from drying out and becoming more challenging to penetrate. If dura proves to be too tough to penetrate, or blunt or multi shank probes are being used, perform a durotomy by lifting the dura with a 27G needle and performing a small incision under saline immersion to prevent the dura from sticking to the brain surface.
4.22. Cover the craniotomy with a hemostatic sponge (see Table of Materials) soaked in cool, sterile saline to protect the dura and brain.

### 5. Surgery: Probe implantation

5.1. Attach the custom microdrive holder (see Table of Materials) to the arm of the stereotactic apparatus. If the microdrive was removed from the microdrive holder after probe preparation, place the microdrive with the attached silicon probe into the microdrive holder. Angle the stereotax arm as required to reach the desired target brain area. Place crown ring with attached amplifier onto the three vertical pins at the rear of the microdrive holder (see Supplementary Figure S1).
5.2. Lower the microdrive to within approximately 0.5 mm of the craniotomy, then use forceps to connect the GND/REF header pin(s) attached to the probe to the corresponding GND/REF pin(s) implanted on the skull (see Steps 4.14-4.15). See Supplementary Figures S3 and S4 for example of drive, craniotomy and GND/REF pin placement.
5.3. Once in place, optionally secure the pin(s) with a drop of conductive silver epoxy (see Table of Materials) for a more robust connection. Once silver epoxy is cured, cover the connected pins in a small amount of dental cement (see Table of Materials) to make sure the connection stays stable over long periods of time, and that there is no electrical connection with the surrounding tissues and/or implant elements.
5.4. Remove the hemostatic sponge from the craniotomy (see step 4.22).
5.5. Position the stereotactic arm with the microdrive over the craniotomy. Note: If the probe is retracted, make sure that the microdrive is placed in a way that the probe would touch down on a part of the craniotomy that does not contain large blood vessels.
5.6. Lower the microdrive, if necessary adjusting in location and angle, until the probe shank touches the dura or brain surface (see step 4.21) in the target area.
5.7. Mix dental cement in the designated ceramic dish (see Step 3.2), and cement the base of the microdrive in place, focusing on the three sides of the microdrive base that are not facing the electrode. Ensure that the cement does not touch the microdrive above the removable ‘base’ (see Figure 1D). Make sure that any space between the base and skull is covered fully with dental cement. Clean the ceramic dish and put it back in the fridge/freezer. Wait for the cement to cure, approximately 10–15 minutes. Note: Leave a small gap between the microdrive base and skull, and use cement in its most fluid form to fill it. Once the cement has thickened slightly, build up the cement between the walls of the microdrive base and the skull. Always use very small amounts of cement, as the flow of the substance can be unpredictable and larger volumes may flow into undesired regions. Small amounts of hemostatic sponge dipped in saline can be used to cover portions of the craniotomy. If cement should accidentally flow onto the craniotomy, remove the cement with forceps once it enters a film-like consistency.
5.8. Lower the silicon probe onto the brain, carefully monitoring probe position through a microscope. When the probe shanks touch the brain, lower the probe quickly by ∼ 250 μm, (one full turn of the screw is 282 μm) to ensure that the probe breaks through the resistance of the dura/cortical surface. Verify this visually. If the probe has not broken into the cortex, wait for 5 minutes, then attempt to etch through the dura with the shank tip by repeatedly raising and lowering the probe by a few tens of micrometers whilst the dura/cortex is under tension from the probe tip.
5.9. Once the probe has broken through the surface of the cortex, gradually lower it at a slower pace (100-200 μm /minute) until either the target coordinates are reached or the probe has moved by more than 1000 μm. If the target requires the probe to move by more than 1000 μm, advance the probe in steps of maximally 1000 μm/session over the following recording sessions until the target coordinates are reached. Note: Skip this step if monitoring neuronal signals while lowering the silicon probe is preferred. Steps for this are described in section 7.
5.10. Prepare silicone elastomer according to instructions (see Table of Materials) and dispense a small drop of it into the craniotomy using a 1mL syringe (see Table of Materials).
5.11. Once dry, cover silicone elastomer with 50/50 mix of bone wax and mineral oil. This step further protects the probe and prevents accumulation of debris and dry plasma over the craniotomy, making extraction simpler and safer. Exercise caution, as working around the probe whilst it is lowered can lead to breakage.

### 6. Surgery: Implantation of Faraday cage

6.1. When the dental cement has fully solidified, loosen the microdrive holder by loosening the lateral screw fixating the drive with an Allen key (see Supplementary Figure S1). Gently retract the holder by approx. 1 cm, so that the microdrive is free-standing, but the probe amplifier/connector remains fixed to the implant holder without stretching the flex-cable.
6.2. Place the pre-made crown and Faraday mesh around the headplate by stretching the cage at the opening and slotting it over the microdrive and Flex-cable horizontally, then fix onto the headplate with dental cement. Note: Make sure to close all spaces between Faraday cage and skull with dental cement to protect the implant from contamination.
6.3. Put the Faraday crown ring (see Table of Materials) with probe connector/headstage over the crown, aligning the integrated holder for the probe amplifier/connector with the area marked by an indented ‘X’ on the Faraday crown(see Step 2.13).
6.4. Secure the ring to the Faraday cage with a small drop of cyanoacrylate glue or dental cement at each spoke-ring junction.
6.5. Once the Faraday ring with integrated probe amplifier/connector is secured in place, fully retract the stereotactic arm with the microdrive holder. See Supplementary Figure S3 for a step-by-step guide on the assembly of these components.

### 7. Post-surgery test recording

7.1. Connect the probe amplifier/connector to the recording hardware and start a recording.
7.2. If the probe has not yet reached its target location during the initial insertion (see step 5.9), slowly turn the microdrive screw counterclockwise to lower the probe while monitoring neuronal signals. Signals should change a) when electrodes touch the layer of silicone elastomer above the craniotomy, and b) when the electrodes begin to move into the brain (see step 7.2). High-frequency neuronal activity will be registered by electrodes that are fully inserted in the brain, while electrodes that are in contact with the CSF on the brain surface will typically show a low-pass-filtered neuronal population signal without spiking activity (akin to an EEG trace), and recording sites in air will register increased electrical noise. Note: It is possible to additionally verify the probe insertion depth of the probe by measuring the impedance of individual channels after the test recording. Channels in contact with air should show high impedance (indicating an open circuit), and impedances like the ones measured before surgery for the channels touching CSF or already in the brain. Advance the silicon probe by a maximum total distance of approximately 1000 µm per session, with a maximum speed of approximately 75 µm per minute (see Step 5.5).
7.3. When neural local field potentials are visible across the probe and/or you have advanced the probe by a maximum of 1000μm, end the test recording and disconnect the head-stage connector.

### 8. Recovery

8.1. Cover the Faraday cage with self-adherent veterinary wrap (see Table of Materials).
8.2. End anesthesia and let the animal recover for a few days following approved experimental guidelines.
8.3. If the electrodes on the silicon probe are not yet at the desired target location, turn the screw of the microdrive in small steps with a maximum of four full turns (or approx. 1000 µm) per session. If necessary, repeat this procedure over several days until the target is reached. Combining the probe movement with simultaneous recordings to evaluate electrophysiological activity in areas transversed is recommended.

### 9. Behavioral experiments and chronic recordings

9.1. For chronic head-fixed recordings during task performance, attach the headplate at the base of the Faraday cage to the head-fixation clamp by manually opening the clamp and clamping the implanted headplate in (see Figure 1C, E and F). Note: If head fixation is not needed, this implant system can also be used for freely moving recordings. For freely moving recordings, skip steps 9.1 and 9.7.
9.2. Remove self-adherent veterinary wrap from implant Note: To minimize discomfort for the animal, we suggest to already start a simple, rewarding behavioral task before this step as a distraction while the experimenter works with the implant.
9.3. Attach amplifier/connector to recording equipment.
9.4. Conduct neuronal recordings as animal performs task. Note: If the goal is to maximize the number of extracellular units recorded, move the shuttle by a few tens of µm whenever the neural yield in a location decreases. Note that after moving the probe, the signal can take minutes to hours to stabilize. Therefore, it might be beneficial to move the probe at the end of a session so that the signal can recover until the start of the next session.
9.5. At the end of behavioral recording, disconnect recording equipment and cover implant in new veterinary wrap.
9.6. Open head-fixation clamp to detach animal from head fixation.

### 10. Probe recovery

10.1. At the end of the final recording, retract the silicon probe as far as possible onto the microdrive by turning the screw clockwise. This can be done while the animal is head-fixed and behaving, or with the animal anaesthetized in the surgical setup. You can chart the exit of the probe from the brain by monitoring neuronal signals simultaneously and checking for the signature of electrodes being immersed in the brain, touching the brain surface, or in contact with air (see Step 7.3). Note: Depending on the histology protocol and probe, perform electrolytic lesions before retracting the probe to determine the exact location of some of the electrodes on the probe. If monitoring the probe exit via neuronal recording is not necessary, it is also possible to retract the probe once the animal has been terminated.
10.2. Terminate the animal following approved guidelines (this includes perfusing the animal if fixating the brain for subsequent histology is planned).
10.3. Wait for ∼ 10 minutes after the animal has died. Then head-fix the animal in the stereotax, making sure that the animal’s head cannot move during explant to prevent probe breakage.
10.4. Apply a drop of saline on top of the craniotomy and let it soak for a few minutes to soften dried biological tissue on the probe shank and lower the chance of shank breakage.
10.5. Place the stereotactic holder approximately 0.5 cm above the microdrive. Then cut the upper end of the spokes of the Faraday cage with small surgical scissors (see Table of Materials) to free up the Faraday ring holding the amplifier/connector and transfer the ring back onto the vertical pins at the top of the stereotactic holder (see step 5.1 and Supplementary Figure S1).
10.6. Carefully cut away the copper mesh with the same surgical scissors by cutting out u-shaped areas of mesh between the spokes of the Faraday crown. Then cut off the crown’s plastic spokes at the base. Note: Avoid bending the printed plastic spokes as you cut them, as they may snap and send plastic debris flying in the direction of the probe.
10.7. Lower the stereotactic holder until the microdrive can be fixated in the holder using the holder’s lateral screw, fixate the microdrive, then loosen the T1 screw that connects the microdrive body to the microdrive base.
10.8. Slowly retract the stereotactic arm with the implant holder to lift the microdrive off its base. Ensure that the microdrive separates from the base at a perpendicular angle (i.e. ‘vertically’ from the base). Note: If the microdrive body and base do not separate easily, verify that the movement of the stereotactic arm is not at an angle compared to the microdrive orientation. If necessary, re-align the holder and microdrive to each other by slightly loosening the fixation of the animal’s head and repositioning it accordingly. Correct alignment is one of the crucial aspects for easy recovery of the microdrive. Also check whether there is any residual dental cement connecting microdrive and microdrive base (see step 5.5). If so, carefully scrape off the cement with a scalpel and/or dental drill depending on the amount of cement.
10.9. Raise the stereotactic arm with the attached probe to create sufficient space below it.
10.10. Remove the animal from the stereotax, and prepare the brain by following an approved histology protocol if desired. Recover the implanted microdrive base and clean it by soaking in acetone for several hours for later re-use.
10.11. Place a clean microdrive base on adhesive putty (see Table of Materials), then lower the microdrive onto the base and tighten the screw. To prevent breakage, monitor the probe position under a microscope throughout the process. This step can be completed at a later time if the implanted microdrive base needs to be cleaned for re-use first. Note: This protocol calls for the use of adhesive putty as a platform for the base, which is vital as it both secures the base whilst also having a degree of give, ensuring the base does not slip and collide with the probe. The putty should be shaped into a vertical ‘cliff face’ on the side of the microdrive base where the probe will be lowered. This ensures that if the probe is lowered past the base, it does not make contact with putty underneath. The putty ‘tower’ should also be tall enough that if it is lowered past the microdrive base, the probe does not make contact with the table surface on which the putty is placed. Finally, secure the putty well to the surface to prevent it from slipping or falling. When lowering the microdrive onto the microdrive base held by the putty, ensure a side profile view fromthe microscope to monitor the progress that as the probe is lowered, it does not collide with either the base or the putty.
10.12. Clean and sterilize the probe following the manufacturer’s instructions. For most commonly available probes, this includes a 12-hour soak in enzymatic cleaner (see Table of Materials), followed by rinsing in demineralized water and sanitization in alcohol. This can be done by lowering the probe into a sufficiently large beaker containing the enzymatic cleaner whilst still attached to the microdrive holder on the stereotactic arm. Note: If desired, measure the impedances of the electrodes on the probe after cleaning to monitor the potential degradation of individual electrodes.
10.13. Store the microdrive with the cleaned probe safely until the next experiment.

## REPRESENTATIVE RESULTS

This protocol presents a chronic implantation system that enables researchers to implement light-weight, cost-effective and safe chronic electrophysiology recordings in behaving mice (Fig. 1). The main factors that determine successful application of this approach include: complete cement coverage of the skull, a minimally invasive and properly protected craniotomy, secure attachment of the microdrive and wiring to the skull and complete continuity of protective Faraday material. When these points are accounted for, high-quality recordings can be reached consistently. Here representative results pertaining to the following main aspects of surgery success are shown:

1. Is the implant interfering with animal behavior or well-being?
2. Is signal quality high, and can signals be maintained over prolonged periods of time?
3. Can recordings be combined easily with task performance?

To assess the impact of the implant on animal behaviour, we analysed tracked locomotion patterns in five implanted animals. Figure 2A shows an example of an animal freely moving inside of a play cage for 10 minutes before and one week after implant. One can see that movement patterns are unchanged. This observation is confirmed by Figures 2B, C showing the distributions of movement speeds and head directions across animals. Both running speed and directions were largely unchanged before and after implantation, and if anything, running speeds seemed to be slightly elevated after surgery. Supplementary Video S1 shows a short video recording of an animal 6 days after implantation surgery. Typical home cage behaviors like locomotion, grooming, rearing and foraging in the home environment are all visible and indicate successful recovery from surgery, as well as general health. The low behavioural impact of the implant is most likely due to its low weight and manageable height.

**Figure 2.**
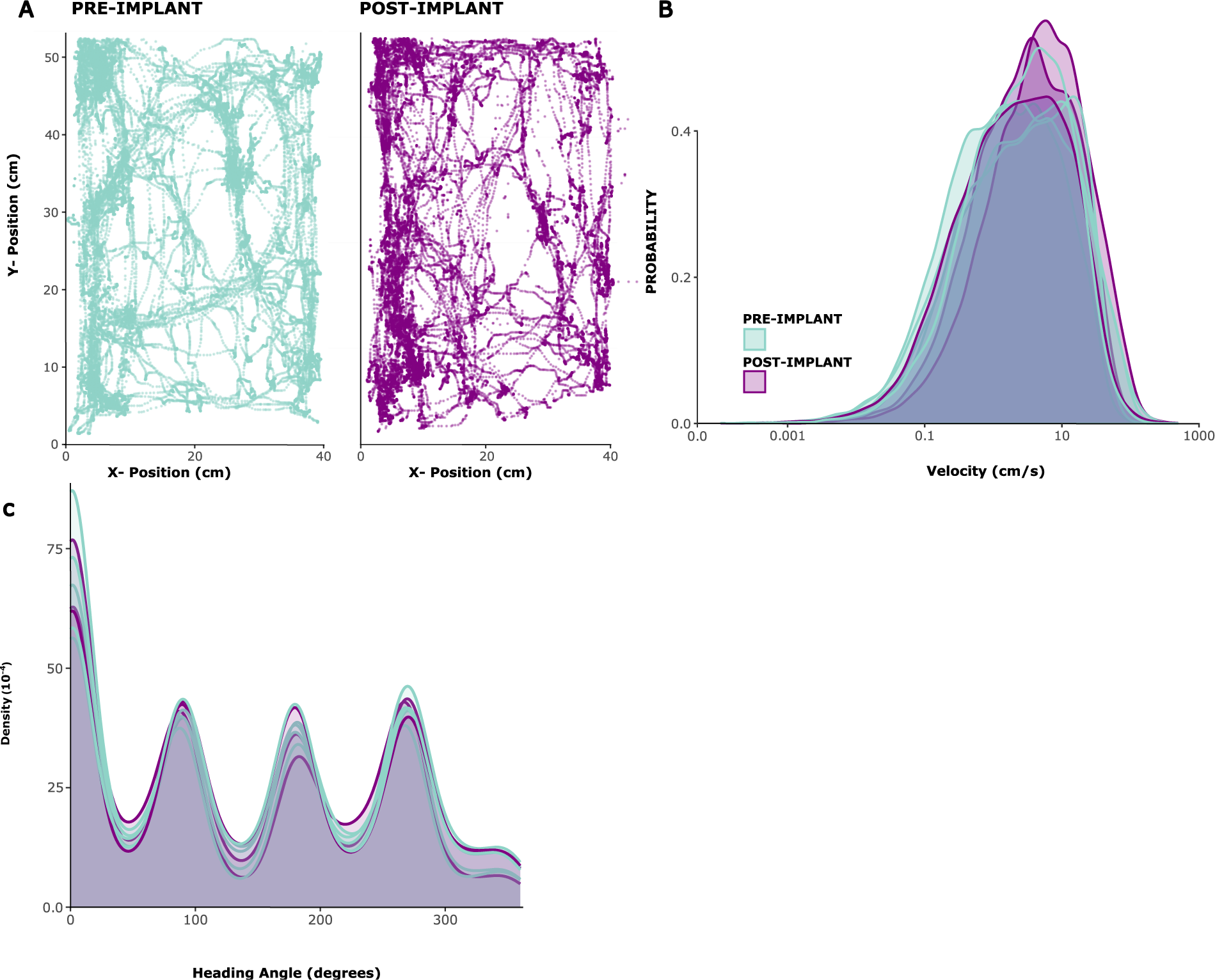
Locomotion before and after surgery. Example locomotion of an animal before (left panel) and after (right panel) implantation. x/y coordinates are in centimeters, points show position of the animal at each timepoint over a period of 10 minutes. (B) Distribution of movement speeds in cm/s for 5 sessions before, and 3 sessions after implantation in 5 animals. (C) Kernel density for probability of movement in different directions, for the same sessions analyzed in (B).

Next, the signal quality in Local Field Potential (LFP) and spiking activity across recording sites is assessed. Here we show representative data from cortical recordings in the primary visual cortex (V1). For validation, putative single unit activity was extracted from broadband neuronal signals recorded in V1 of an awake mouse using Kilosort 3 (see Figure 3). Figure 3A shows the location of extracted single units on the probe shank, Figure 3B shows the corresponding spike wave forms and Figure 3C shows the spiking responses of the same neurons to a Current Source Density (CSD) protocol. In this paradigm, wide-field flashes were presented with a duration of 300ms, at a frequency of 1 Hz (i.e. 300ms on, 700ms off) over 200 trials. Finally, Figure 3D show the same units’ responses to a visual receptive field mapping protocol, consisting of 2000 frames of randomly selected black and white squares on a grey background, each presented for 16.6ms. Squares covered 12 degrees of visual angle each and were selected from a field of 15×5 possible locations, so that the mapping paradigm covered a visual space of -90 to +90 degrees azimuth and -30 to +40 degrees elevation in total. Firing rate responses to each stimulus frame were extracted by analyzing the maximum firing rate across a 16.6ms window, subject to a delay of between 40-140ms identified as optimal per channel based on the maximum activity in each window. This type of recording can be used to guide adjustment of the insertion depth of each electrode, and to assess signal quality after the implant surgery.

**Figure 3.**
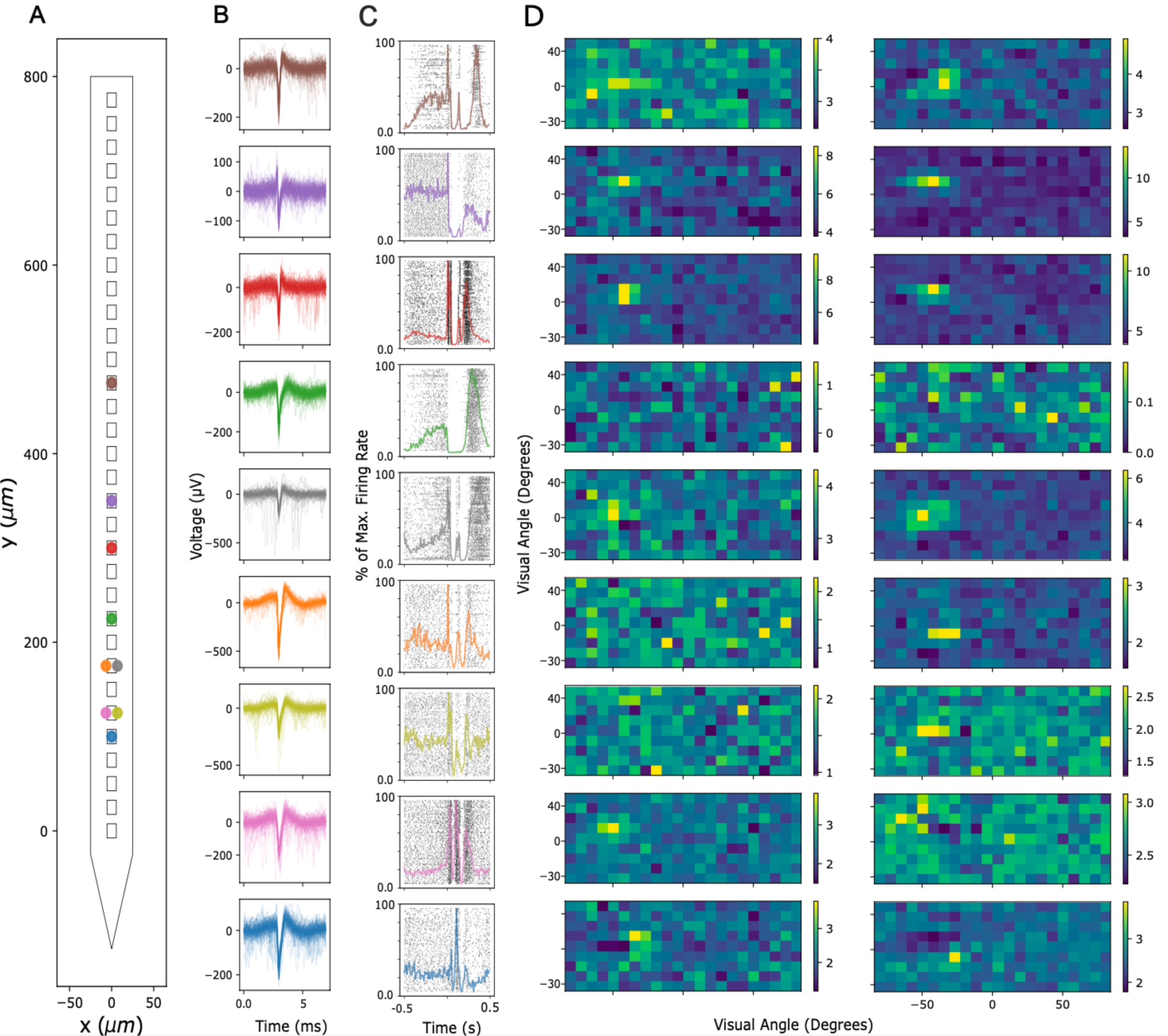
Recorded neuronal signals. (A) Inferred location of single units sorted by Kilosort 3 spike sorting package along the probe’s electrode contacts. (B) Spike waveforms for the same units shown in (A) across 5ms of time. Thin lines: Individual spike waveforms. Thick lines: Average spike waveform. (C) Raster plot of spikes in response to a Current Source Density (CSD) paradigm presenting 300ms wide-field flashes followed by a 700ms black screen. Responses are shown for the same units as in (A) and (B). Superimposed colored lines represent peri-stimulus time histograms (PSTHs) of the same responses. Firing rates for the PSTHs were calculated in 10ms bins and then normalized by the maximum firing rate across the entire PSTH. Time 0 is centered around the widefield flash stimulus. (D) Estimated receptive fields of the same units as in (A) - (C), measured by a Sparse Noise Receptive Field Mapping paradigm. Each plot shows average firing rate activity over a 16.6ms analysis window in response to the onset (left panel) or offset (right panel) of white and black square stimuli. Stimuli were presented for the duration of 16.6ms, located randomly across a 5×15 square grid spanning 180 degree of visual angle horizontally, and 70 degree of visual angle vertically. Firing rate activity was z-scored across the entire receptive field grid (see color bar).

Recording quality remained high across repeated recordings for weeks to months. Figure 4A shows longitudinal LFP recordings from one animal over 15 weeks. LFPs were recorded in response to the CSD paradigm described above (see Fig. 3A-C). Figure 4A shows averaged LFP responses 500 milliseconds following flash onset. In this example, we used a linear probe with 32 channels, with an interelectrode distance of 25 µm. Note that on day 18 the probe depth was adjusted, shifting the probe downwards by 600 µm. Both before and after this adjustment, LFP signals remained stable across recording days.

**Figure 4.**
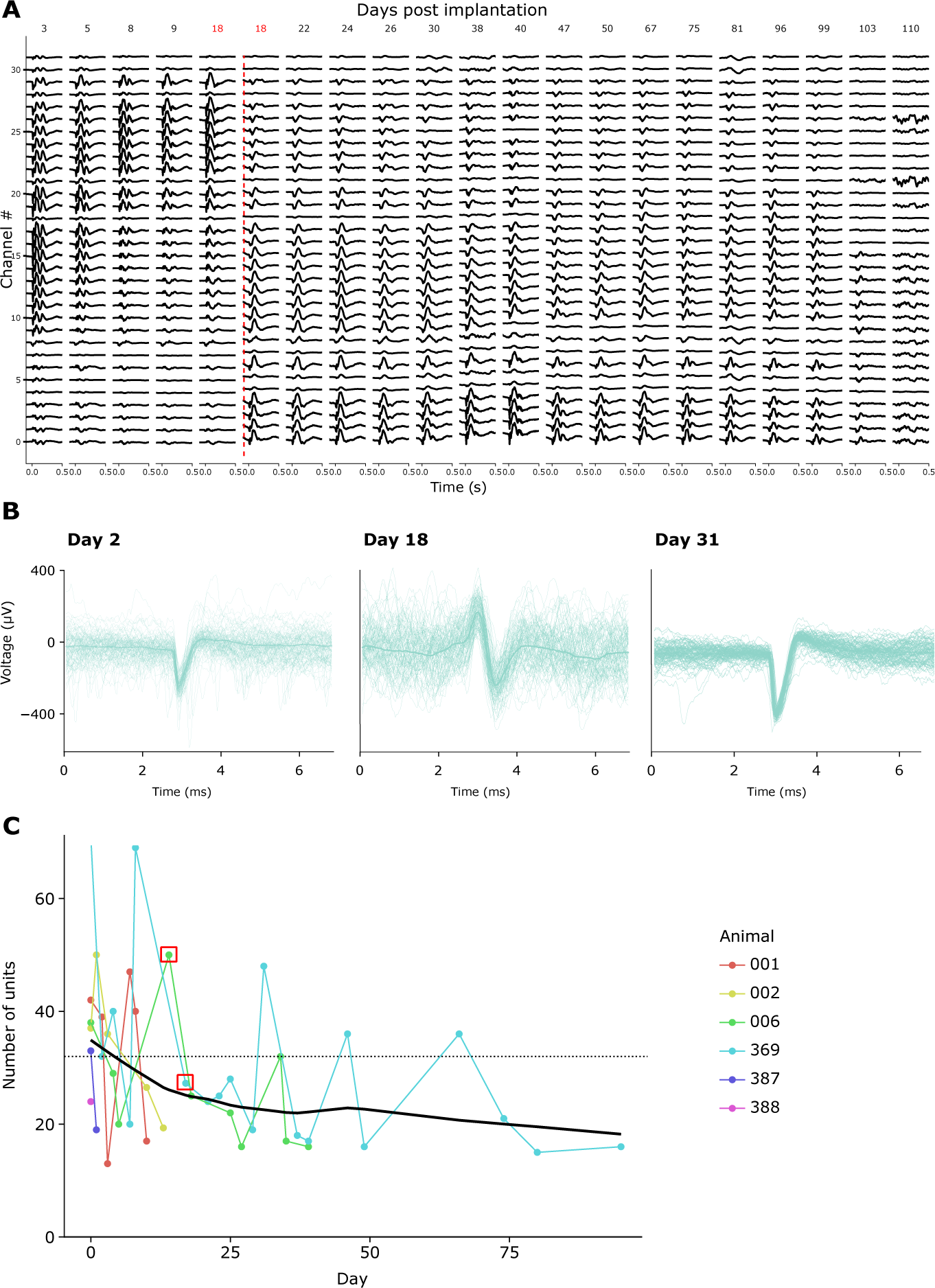
Stability of neuronal recordings over time. (A) Average LFP activity in response to a wide-field flash CSD stimulus, shown across all 32 channels of a chronically implanted probe from 3 to 110 days post implant. Red vertical line denotes probe being lowered to a new location due to channels 0-8 recording from outside the brain by Day 18 post-surgery. (B) Spike waveforms of three example units from the same chronic implant recorded repeatedly across four weeks. Thin lines: Individual spike waveforms. Thick superimposed line: Average spike waveform. (C) Number of putative single units detected by Kilosort 3 across recording days for six animals (see inset legend). Red Square denotes days when probe was moved. Dotted line denotes number of electrodes per implant used in these recordings (32).

Consistent with this, spike waveforms of putative single units were discernible over many recordings. Figure 4B shows representative example spike waveforms from three recording sessions across a month of recordings, demonstrating that single unit activity can be identified successfully over time. Figure 4C shows the overall number of putative single units extracted from chronic recordings in six animals, spanning a window of up to 100 days. Single units were defined according to the default criteria of kilosort 3.0 (see Supplementary Table S1). As one can see, the number of clearly defined single units typically amounted to approx. 40 in the first week post-implantation, and then dropped off gradually, moving towards an apparently stable asymptote of approx. 20 units. Given that these recordings were conducted using linear 32-channel probes, this equates to an expected yield of about 1.25 single units per electrode directly after implantation, declining to approx. 0.65 single units per electrode in long-term recordings. Repeated connection to the implant’s amplifier/connector over sessions did not appear to impact either recording quality or implant stability, since the Faraday crown that holds the amplifier/connector can withstand repeated forces of over 10 Newton, an order of magnitude larger than even the maximal mating forces required by standard connectors (see Video S2).

Finally, by providing a modular system including a microdrive as well as a wearable Faraday cage and a headplate that doubles as an implant base and a device for head-fixation, this protocol enables the integration of chronic electrophysiology with head-fixed behavior. Here example data from mice traversing a virtual environment on a spherical treadmill are shown. Figure 5A shows running-related spiking activity of 20 units in an example trial, and Figure 5B shows the diverse but robust relationships between running speed and spiking activity of individual spike-sorted units, as well as a population average for the same effect in Figure 5C, confirming the well-established effect of locomotor activity on neuronal activity in rodent V1 5.

**Figure 5.**
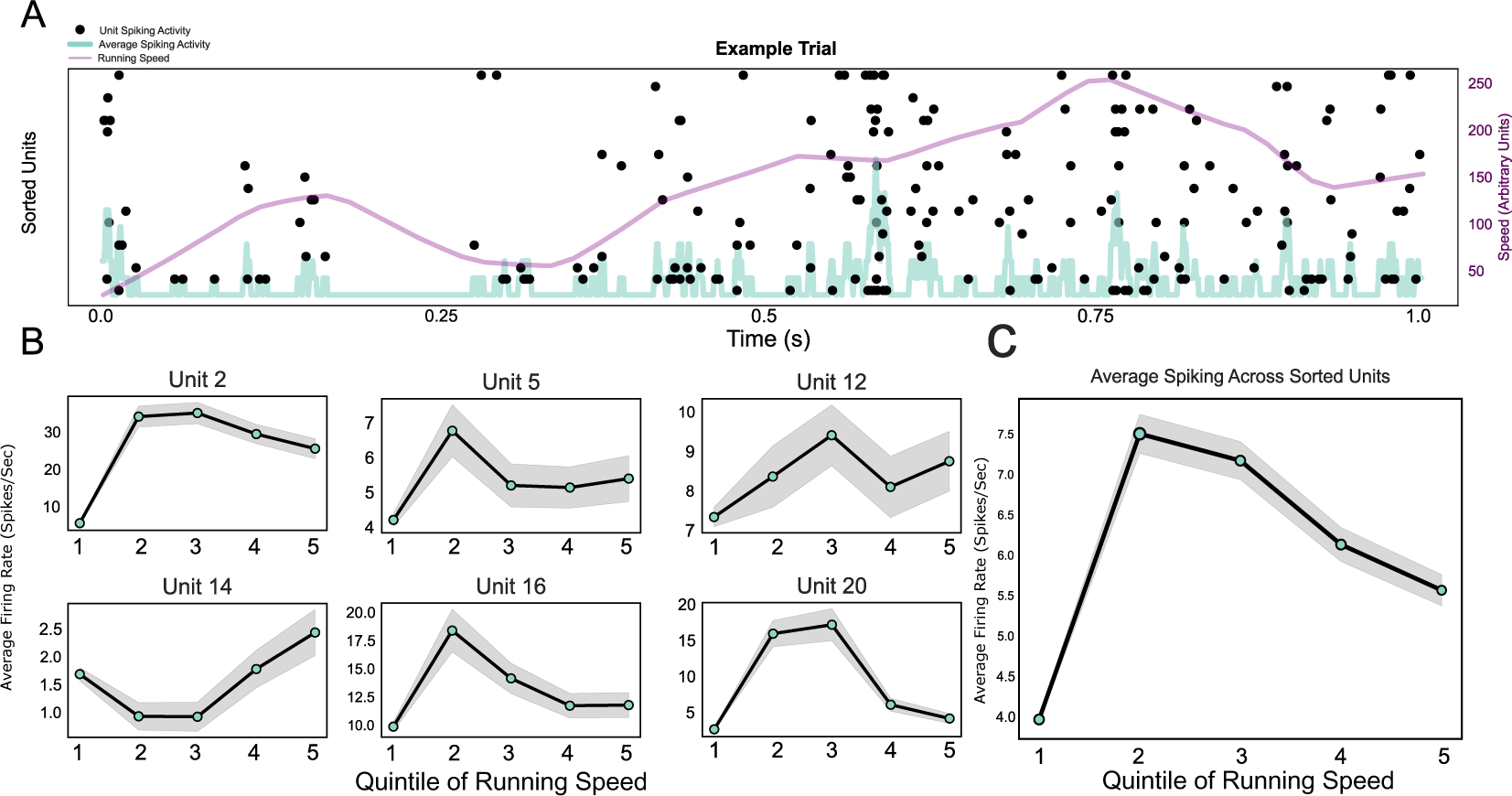
Neuronal responses during head-fixed behavior. (A) Raster plot of single unit responses across an example trial, with running speed (purple line) and average firing rates across all single units (light blue line) superimposed. (B) Single unit activity during different running speed categories, shown for six example units. (C) Average spiking activity across all single units in one example session, plotted across the five quinitiles of the running speed distribution. Running speeds in this session ranged from 0 to 0.88 meters/second.

## DISCUSSION

This manuscript presents a protocol for the fast, safe and standardized implantation of probes, which also allows probe recovery and re-use at the end of the experiment. The approach makes use of a modular system of implant components, specifically a microdrive, which is compatible with all common silicon probes and recording systems, a headplate that can be used for head-fixed behavioral experiments, and a wearable Faraday cage to protect the implant. This constellation allows users to flexibly adapt their implant to different experimental paradigms, such as head-fixed versus freely moving behavior or implant miniaturization (without Faraday cage) versus increased long-term signal robustness (with Faraday cage) - without having to sacrifice the standardization of the implant in the process.

This approach makes chronic electrophysiological recordings more standardized (through prefabricated elements that do not require assembly by hand), less costly (through probe recovery), less time-consuming (by simplifying surgery steps) and more easily compatible with animal welfare and behavior (through decreased implant size and stress-free head fixation). As such, this protocol aims to make electrophysiological implants in behaving rodents attainable for a broader range of researchers, beyond the pioneering labs at the cutting edge of the field.

To achieve this aim, protocol presented here minimizes the trade-off between several often equally crucial aspects of microdrive implants, namely flexibility, modularity, ease of implantation, stability, overall cost, compatibility with behaviour, and probe reusability (see Table 1). Currently available approaches often excel at some of these aspects, but at a steep cost to other features. For instance, for use-cases that demand absolute implant stability over long time periods, the best implant approach may be to directly cement the probe onto the skull^25^. However, this also prevents probe re-use, as well as repositioning of recording sites in case of bad recording quality, and it is incompatible with standardized implant placement. Similarly, while e.g. the AMIE drive provides a lightweight, low-cost solution for recoverable implantation of probes, it is limited to single probes and restricted in the placement of the target coordinates17. At the opposite end of the spectrum, some commercially available nano-drives (see Table 1) are extremely small, can be placed freely on the skull and maximize the number of probes that can be implanted on a single animal16. However, they are expensive compared to other solutions, require experimenters to be highly skilled for successful implant surgeries, and prohibit probe re-use. The microdrive developed by Vöröslakos et al.21, a light-weight version of which is also part of this protocol, sacrifices small implant size for better ease-of-use, lower price, and probe re-usability.

**Table 1.** Comparison of popular strategies for chronic probe implants in rodents. Availability: whether the microdrive is open source (for researchers to build themselves), commercially available, or both. Modularity: Integrated systems consist of one or few components that are in a fixed relation to each other, while modular systems allow free placement of the probe / microdrive relative to the protection (head gear/Faraday cage) after production of the implant (e.g., at time of surgery). Modularity was determined from published information or implantation protocols of the listed implants. Headfix: Yes: The implant has mechanisms for head-fixation integrated in its design, X: The implant leaves the space to add an extra headplate for fixation without big issues, No: The design of the implant likely creates space issues or requires substantial design modifications for use with head fixation. Probe placement: Restricted: Probe location is limited at the implant design stage. Flexible: Probe location can be adjusted even during surgery. Number of probes: the number of probes that could be implanted. Note that implanting >2 probes on a mouse does pose a significant challenge independent of the chosen implant system. Probe reusability: yes, if the probes can, in theory, be reused. Weight/size: weight and bulkiness of the implant.

| Implant | Availability | Modularity | Headfix | Electrode placement | Number of probes | Probe reusability | Weight |
| --- | --- | --- | --- | --- | --- | --- | --- |
| DREAM | Open source | Modular | Yes | Flexible | 1-2 | Yes | 1.9g |
| AMIE<sup>17</sup> | Open source | Integrated | Yes | Restricted | 1 | Yes | 1.5g |
| Chung<sup>26</sup> | Open source | Modular | Yes | Flexible | 2 | Yes | 3.34g |
| Vöröslakos<sup>21</sup> | Open source | Modular | No | Flexible | 2-3 | Yes | 2.2g |
| Apollo drive<sup>27</sup> | Open Source | Integrated | No | Restricted | 2 | Yes | 2.29g-1.26g |
| Jones drive<sup>28</sup> | Open Source | Integrated | Yes | Restricted | 2 | Yes | <3g (1 probe), <4g (2 probes) (both w/ probe & cement) |
| NeuroNexus<sup>29</sup> | Commercial | Integrated | No | Restricted | 1 | No | 0.85g |
| NanoDrive<sup>16</sup> | Commercial | Modular | X | Flexible | 2-3 | No | 0.6g |
| Van Daal<sup>30</sup> | Open source | Integrated | No | Restricted | 2 | Yes | 3.5-6.8g Incl. Probe + Headstage |
| Probe directly cemented on skull | Open | n/a | X | Very flexible | 2-3 | No | 0g |

To create a system that reconciles these different requirements more seamlessly, the DREAM implant was designed on the basis of the Vöröslakos implant 21, but with several fundamental modifications. First, to reduce overall implant weight, the microdrive used here is produced in machined aluminium rather than 3D-printed stainless steel, and the Faraday crown is miniaturized, achieving an overall weight reduction of 1.2 - 1.4g depending on the choice of headplate material (see Table 2). Second, the headplate surrounding the microdrive was designed to allow for an integrated head fixation mechanism that enables fast and stress-free head fixation while doubling as a base for the Faraday cage, giving access to most potential target areas for neuronal recordings, and adding only minimal weight to the implant. The flat shape of the fixation mechanism and lack of protrusions also ensures minimal impairment of animals’ visual field or locomotion (see Fig. 2A-C), a clear improvement over previous systems^31, 32^. The Faraday crown and ring that are fixed onto the headplate were also substantially altered compared to previous designs. They now do not require any ad-hoc adaptation (e.g. in terms of connector placement) or soldering throughout the surgery, removing potential causes of implant damage and of unpredictable variance in implant quality. Instead, the DREAM implant provides multiple standardized crown ring variations that allow to place each connector at one of four pre-defined positions, minimizing variability and effort during surgery. Finally, by optimizing the implant system for probe recovery, the DREAM implant allows experimenters to drastically cut the cost as well as preparation time per implant, since microdrive and probe can typically be recovered, cleaned and re-used together.

**Table 2.** Comparison of component weights between the DREAM implant and the implant described by Vöröslakos et al. (2021).

| Implant Component | Schröder, Taylor et al., | Vöröslakos et al., |
| --- | --- | --- |
| Drive | 0.5g | 0.87g |
| Faraday Cage (mesh+support) | 1.0g | 1.96g |
| Headplate | 0.2-0.5g | N/A |
| Base | N/A | 0.19g |
| Screws | N/A | 0.15g |
| Total | 1.7-2.0g | 3.17g |

For a more exhaustive overview of the trade-offs posed by different implant systems, see Table 1. While the approach presented here does generally not provide maximal performance compared to all other strategies e.g. in terms of size, stability or cost, it operates in the upper range across all these parameters, making it more easily applicable to a wide range of experiments.

Three aspects of the protocol are particularly crucial to adapt to each specific use case: The constellation of ground and reference, the technique for cementing the microdrive, and implant validation via neuronal recording. First, when implanting the ground and reference pins, the goal was to identify the sweet spot between mechanical/electrical stability and invasiveness. While e.g. floating silver wires embedded in agar are less invasive than bone screws^33^, they are likely more prone to becoming dislodged over time. The use of pins, coupled with agar, ensures a stable electrical connection, whilst also having the advantage of being easier to control during insertion, avoiding tissue trauma. Ground pins cemented to the skull are unlikely to become dislodged, and in the event of the wire becoming separated from the pin, reattachment is usually simple due to the larger surface area and stability of the implanted pin.

Second, cementing of the microdrive should generally occur prior to insertion of the probe in the brain. This prevents lateral movement of the probe inside the brain if the microdrive is not perfectly fixed in the stereotactic holder during insertion. To check the placement of the probe before cementing the microdrive in place, one can briefly lower the tip of the probe shank to ascertain where it will contact the brain, since extrapolating the touchdown position can be difficult given the microscope’s parallax shift. Once the microdrive position is established, one optionally can protect the craniotomy with silicone elastomer prior to cementing the microdrive to ensure that the cement does not accidentally make contact with the craniotomy, however lowering the probe through the silicone elastomer is not recommended, as silicone elastomer residue can be pulled into the brain and cause inflammation and gliosis.

Third, depending on the experimental protocol used, a test recording directly after surgery may or may not be useful. Largely, neuronal activity recorded right after probe insertion will not be directly representative of activity recorded chronically, due to factors like transient brain swelling and tissue movement around the probe, meaning that both insertion depth as well as spike waveforms are unlikely to stabilize directly. As such, immediate recordings can mainly serve to ascertain general signal quality and implant integrity. It is recommended to utilize the moveable microdrive sled in subsequent days post-surgery once the brain has stabilized to fine-tune the position. This also helps to avoid moving the probe by more than 1000µm per day, minimizing damage to the recording site, and thus improving recording site longevity.

Finally, users may wish to adapt the system to record from more than one target location. As this system is modular, the user has a lot of leeway on how to assemble and place components in relation to each other (see above and Supp. Figs. S3 and S4). This includes modifications that would allow a horizontally extended shuttle to be mounted on the microdrive, allowing for multiple probes or large multi shank probes to be implanted, as well as the implantation of multiple individual microdrives (see Supp. Figs. S3-S4). Such modifications only require the use of an adapted crown ring, with an increased number of mounting zones for connectors/interface boards/headstages. However, the space limitations of this design are dictated by the animal model, in this case the mouse, which makes stacking multiple probes onto one microdrive more attractive in terms of footprint than implanting several microdrives independently of each other. The microdrives used here can support stacked probes, and thus the only real limitation is the number of headstages or connectors that can fit the space and weight constraints defined by the animal model. Spacers can also be used that further increase non-vertical mounting and insertion paths.

In conclusion, this protocol allows for inexpensive, lightweight and importantly adjustable implantation of a probe, with the added benefit of a microdrive design that prioritizes probe recovery. This tackles the problems of prohibitive costs of single-use probes, the high barrier of surgical and implantation skill, as well as the fact that commercial solutions for chronic implantation are often difficult to adapt to unique use cases. These issues pose a pain point to labs already using acute electrophysiology, and a deterrent to those that do not yet undertake electrophysiology experiments. This system aims to facilitate the wider uptake of chronic electrophysiology research beyond these limitations.

## ACKNOWLEDGMENTS

This work was supported by the Dutch Research Council (NWO; Crossover Program 17619 “INTENSE”, TS) and has received funding from the European Union’s Seventh Framework Program (FP7/2007-2013) under grant agreement No. 600925 (Neuroseeker, TS, FB, PT), as well as from the Max Planck Society.

