## Supplemental Figures 1-4 for "The DREAM implant: A Lightweight, Modular and Cost-Effective Implant System for Chronic Electrophysiology in Head-fixed and Freely Behaving Mice"

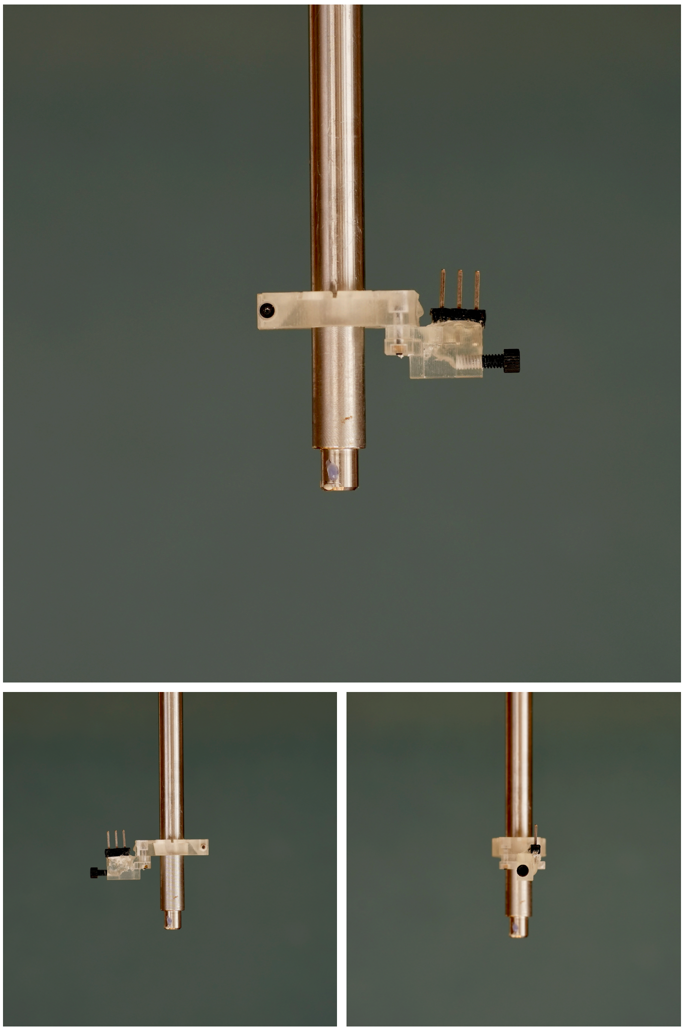

***Supplementary Figure S1.*** *Figure showing images of drive holder. Printable design files can be found in the corresponding Github repository (https://github.com/zero-noise-lab/dream-implant/).*

A B

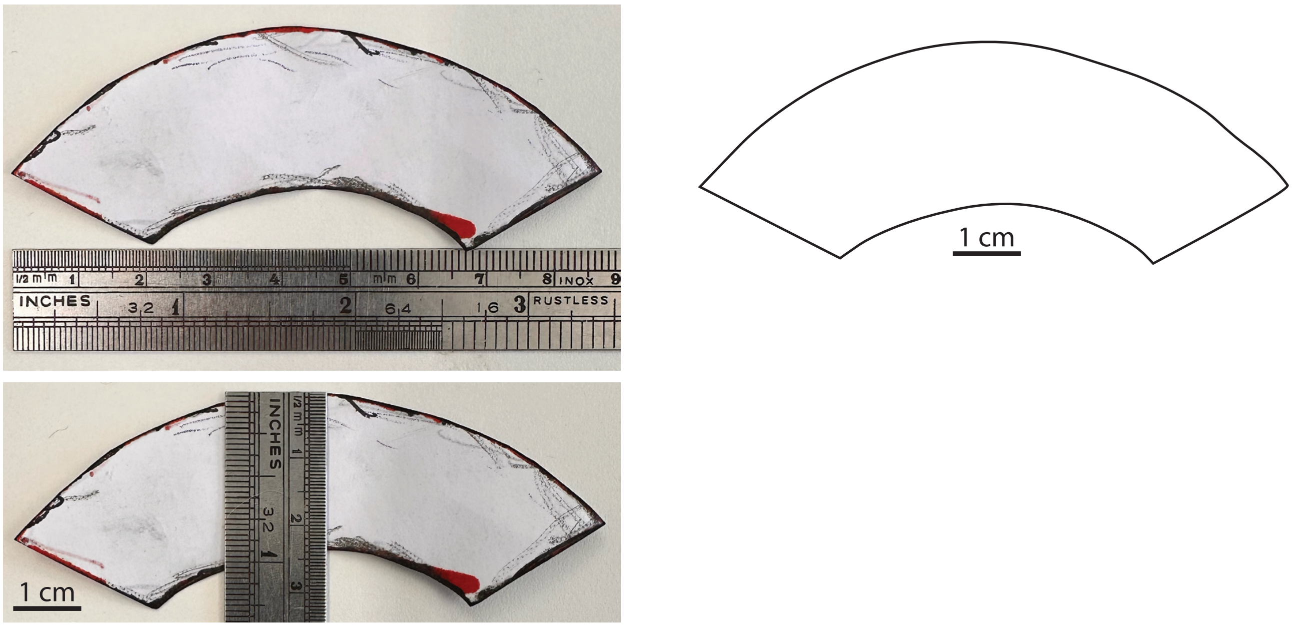


***Supplementary Figure S2.*** *Template for copper mesh. Print the template with original scaling and use the stencil for cutting out the copper mesh (Step 2.12). Use the scale bar for verifying and, if necessary, adjusting scaling of the print.*

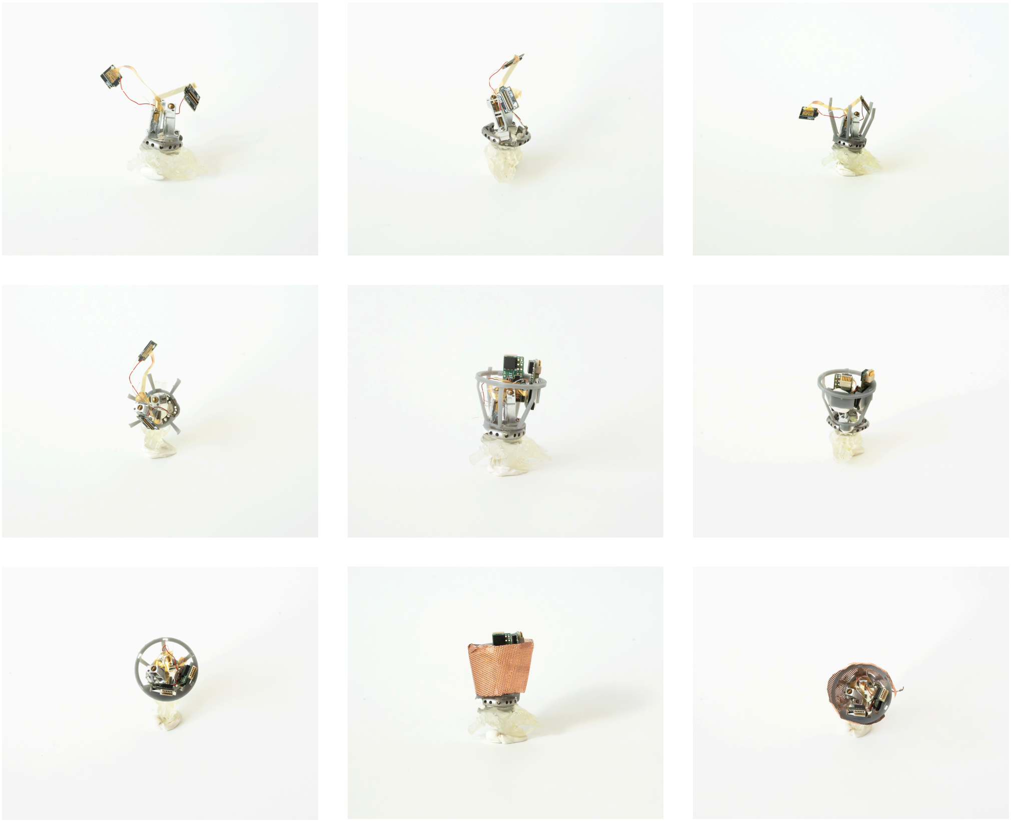

***Supplementary Figure S3.*** *Photo series showing the assembly steps of the implant during surgery. Two microdrives as well as two amplifiers are installed in this case.*

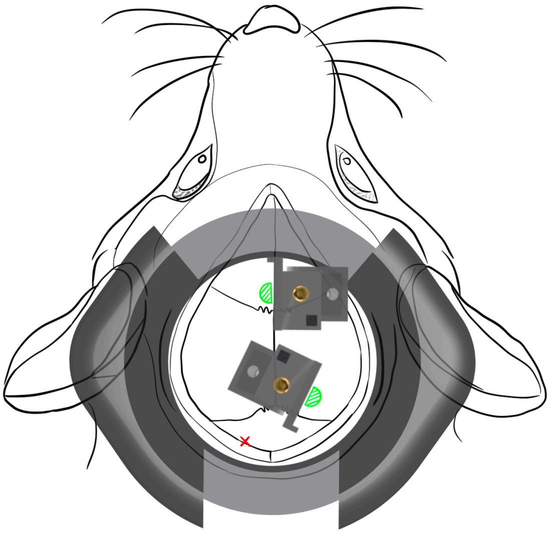

***Supplementary Figure S4.*** *Drawing of mouse skull featuring example placement of drives, craniotomies (in green) and GND/REF pin (in red). Pin location is suggested due to placement in cerebellum, unlikely to interfere with cortical recordings.*

***Supplementary Table S1.*** *Table showing default parameters used by Kilosort 3 when identifying single units in the recordings shown in Figure 3, 4 and 5.*

***Video S1.*** *Video showing animal locomotor activity post implant. Video taken after 5 day recovery phase is complete, showing normal locomotor behaviour, as well as adaption to the size and weight of the implant. Animal can be seen normally exploring a play cage containing environmental enrichment.*

***Video S2.*** *Video showing force being applied onto the assembled Faraday crown. The forces withstood by the Faraday crown are approximately one order of magnitude larger than the connection force needed for standard connectors such as 4-pin polarized nano connectors.*
